# Rapamycin Reverses the Hepatic Response to Diet-Induced Metabolic Stress That Is Amplified by Aging

**DOI:** 10.1101/2025.07.25.666837

**Authors:** Aaron Havas, Adarsh Rajesh, Xue Lei, Jessica Proulx, Karl N. Miller, Adam Field, Andrew Davis, Marcos Garcia Teneche, Armin Gandhi, Peter D. Adams

## Abstract

Aging is associated with increased susceptibility to metabolic stress and chronic liver disease, yet the interactions between age and metabolic stressors and the potential for ameliorating interventions remain incompletely understood. Here, we examined the hepatic response of young (7-month-old) and old (25-month-old) C57BL/6 male mice to a 9-week high-fat diet (HFD) and assessed whether rapamycin, a well-established pro-longevity intervention, could mitigate age-exacerbated effects. While both age groups developed metabolic-associated steatohepatitis (MASH), older mice displayed more severe hepatic steatosis, inflammation, and transcriptional dysregulation. Transcriptomic profiling of whole livers and purified hepatocytes revealed that aging amplifies HFD-induced inflammatory and metabolic gene expression changes, including activation of immune pathways and suppression of metabolic pathways. Notably, treatment of aging mice with rapamycin reversed the majority of HFD-driven transcriptional alterations, including upregulation of pro-inflammatory regulators such as Stat1, and dysregulation of metabolic gene networks. Rapamycin also reduced hepatosteatosis, total body weight, and a tumorigenic transcriptomic signature associated with hepatocellular carcinoma risk. These findings demonstrate that aging intensifies hepatic sensitivity to metabolic stress and identify rapamycin as a promising therapeutic to counteract age-related liver dysfunction and MAFLD progression.

## Introduction

As the liver ages, it exhibits structural and functional changes including reduced volume, diminished blood flow, and mild fibrosis^1,2^. Immune function becomes dysregulated, with increased immune cell infiltration. Aged liver also exhibits mitochondrial dysfunction, altered insulin signaling and disrupted metabolic homeostasis^3–6^. Together, these changes reduce the liver’s resilience to injury and disease, underscoring its vulnerability in aging ^7,8^.

Metabolic-associated fatty liver disease (MAFLD) is a progressive disease, including liver steatosis. MAFLD is a growing public health burden through its association with cirrhosis and liver cancer development ^9,10^. Obesity and associated metabolic dysfunctions fueled by high-fat diets (HFD) and sedentary lifestyles have become epidemic across all age groups^11^. Obesity increases the risk of MAFLD.

Advanced age exacerbates metabolic dysfunction through altered immune responses, hepatocyte physiology, and diminished regenerative capacity^12^. Older individuals demonstrate altered fat distribution, increased susceptibility to severe steatosis, and greater risk of MAFLD progression, hepatocellular carcinoma^13^ and more severe and treatment-resistant disease^14–16^.

However, while obesity- and metabolic dysfunction-related pathologies have been extensively modeled, most studies examine aging and metabolic stress in isolation, failing to capture their intersection. There remains a critical need to understand how aging modifies the transcriptional and physiological landscape under conditions of metabolic stress, and to identify interventions capable of mitigating these compounded effects.

Rapamycin, a selective inhibitor of mTORC1, has emerged as one of the most promising pharmacologic candidates for promoting healthy aging and extending lifespan^17^ ^18^. Across a wide range of model organisms, rapamycin has been shown to robustly increase longevity, even when administered later in life^5,17,19^. Its pro-longevity effects are attributed to its ability to modulate key aging-related processes, including altered growth signaling, enhanced autophagy, suppression of chronic inflammation, and improved mitochondrial and metabolic function^20^. In metabolic disease models, rapamycin reduces adiposity, improves insulin sensitivity, and protects against hepatic steatosis and dyslipidemia^21–23^. These effects suggest a potent role in mitigating the consequences of metabolic stress. Despite these advances, the ability of rapamycin to counteract physiological and molecular responses to metabolic challenges in aged organisms, particularly in the context of high-fat diet (HFD) exposure, remains incompletely understood^23^.

In this study, we interrogated how aging modifies hepatic responses to metabolic stress in the form of HFD, and whether rapamycin can attenuate these effects. Using a comparative model of young and old mice subjected to HFD, we show that aging increases hepatic steatosis and inflammation, with HFD further suppressing metabolic pathways and activating inflammatory pathways, such that older mice displayed more severe HFD-induced hepatic steatosis, inflammation, and transcriptional dysregulation. Rapamycin treatment reversed these deleterious HFD-induced transcriptional changes, reduced body and liver mass, and improved hepatic histopathology in aged mice. These findings suggest that rapamycin can counteract the exaggerated transcriptional and physiological consequences of metabolic stress in aging, expanding our understanding of its therapeutic potential for age-related metabolic disease.

## Results

To investigate how aging impacts the response to metabolic stress, we examined the effects of a high-fat diet (HFD) on young (5-month-old) and old (22-month-old) C57BL/6 male mice over a 9-week feeding period. Both age groups exhibited comparable weight gain on HFD, with no significant differences in body weight between age groups at collection (Fig. 1A). However, histological analysis revealed notable age-dependent differences: while both young and old mice on HFD developed signs of hepatic steatosis and metabolic-associated steatohepatitis (MASH), as evidenced by fat accumulation (Fig. 1B, green arrows) and hepatocyte ballooning (Fig. 1B, yellow arrows), these features were more pronounced in the older cohort. Additionally, old mice displayed elevated immune cell infiltration^4,24–28^ (Fig. 1B, red arrows).

**Figure 1.**
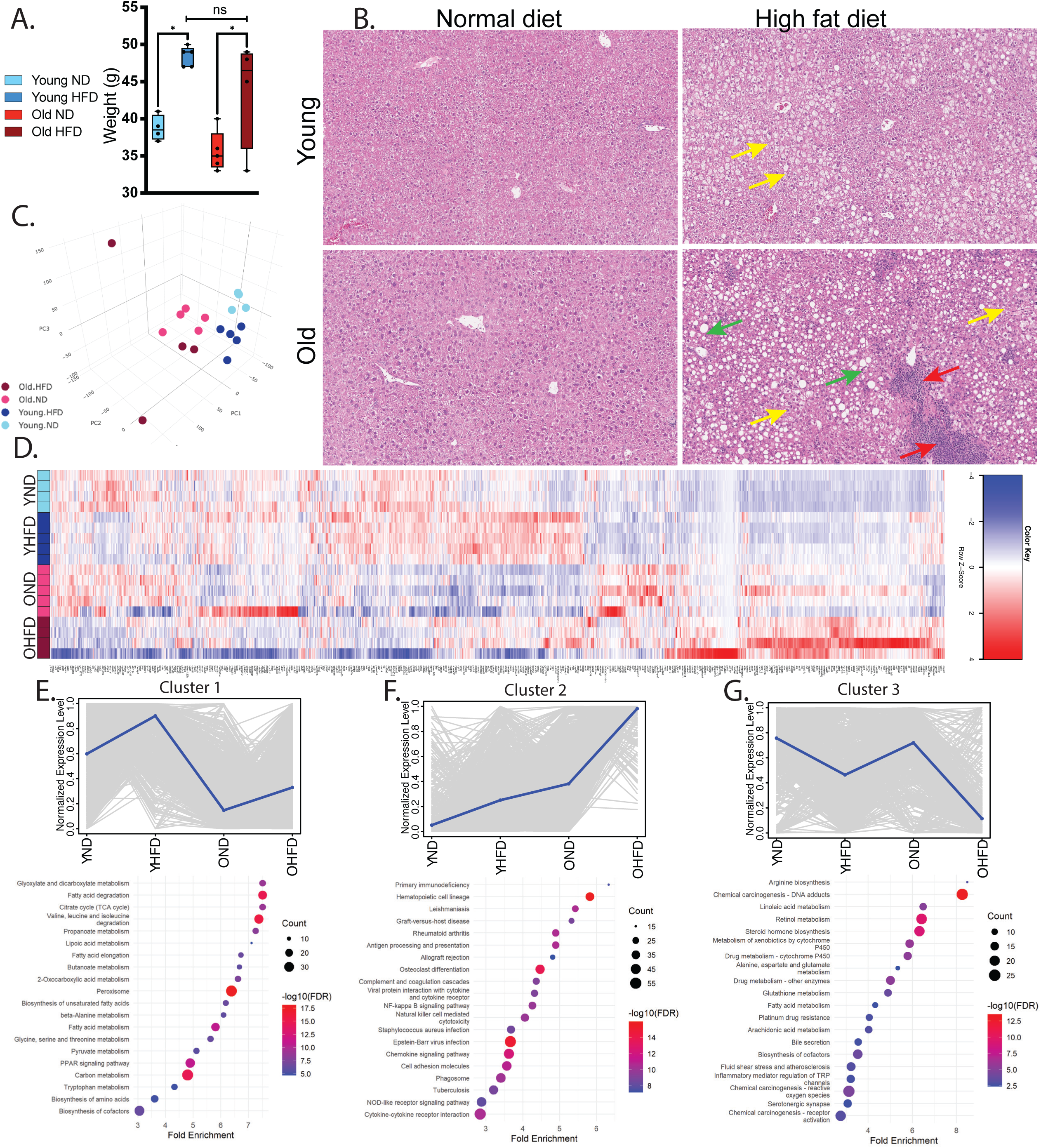
Aging exacerbates high-fat diet (HFD)-induced metabolic stress responses. (A) Body weight distribution (grams) of young (5-month-old) and aged (22-month-old) mice after 9 weeks of HFD or control diet. (B) Representative H&E-stained liver sections. Green arrows indicate fat accumulation; yellow arrows indicate hepatocyte ballooning and red arrows indicate immune infiltration. (C) Principal component analysis (PCA) of whole-liver RNA-seq data. (D) Heatmap showing all differentially expressed genes (DEGs). (E–G) Gene clustering analysis (top panels) with associated enriched KEGG pathways (bottom panels). Statistical analysis of body weights was performed using one-way ANOVA with post hoc Tukey’s test; *p* < 0.05 is indicated by an asterisk (*).

To investigate how aging influences the hepatic response to high-fat diet (HFD) at the molecular level, we performed RNA sequencing (RNA-seq) on whole liver samples. Principal component analysis (PCA) revealed distinct clustering of all groups, albeit with the highest variability observed in the old HFD cohort, suggesting an age-dependent divergence in transcriptional responses to HFD (Fig. 1C). Differential gene expression analysis identified over 4,000 differentially expressed genes (DEGs) across any cohort, with the most substantial difference—3,954 DEGs—between young ND and old HFD groups (Fig. 1D). Clustering of DEGs revealed three major expression trajectories. Cluster 1, encompassing 1,190 transcripts upregulated by HFD in both age groups, was enriched for KEGG metabolic pathways, including fatty acid degradation and peroxisome and carbon metabolism. Cluster 2, comprising 1,244 transcripts elevated with age and further amplified by HFD, was enriched for inflammatory and immune activation pathways such as Graft-versus-host disease and infection responses. Cluster 3, containing 627 transcripts downregulated by HFD in both age groups, included genes involved in metabolism, cytochrome P450, and steroid hormone biosynthesis. These data suggest that while the transcriptional response to HFD shares core similarities between young and old livers, aging amplifies some inflammatory and metabolic changes as identified in clusters 2 and 3, underscoring the heightened vulnerability of aged livers to metabolic stress, also observed at the histological level.

Hepatocytes are the major drivers of liver metabolic functions and exhibit ballooning and fat accumulation in response to HFD (Fig. 1B). Therefore, to achieve a hepatocyte-specific understanding of transcriptional responses to HFD, we performed RNA-seq on isolated hepatocytes from an additional cohort of mice subjected to the same experimental treatment. PCA again revealed the greatest transcriptional divergence between young normal diet and old HFD cohorts, supported by a heatmap showing 5,565 DEGs between any pairwise comparison (Fig. 2A, B). Analysis identified three major gene clusters comprising over 50% of the DEGs (Fig. 2C). Cluster 1, containing 439 transcripts upregulated by HFD in hepatocytes of both young and old mice, was enriched for metabolic pathways, such as fatty acid degradation, peroxisome signaling, and fat digestion, similar to Cluster 1 from whole-liver analysis (Fig. 2C, D). Cluster 2 included 1,033 transcripts elevated with age and further amplified by HFD and was enriched for pathways involved in infection and antigen presentation, again similar to Cluster 2 from whole-liver analysis (Fig. 2C, E). Cluster 3, comprising 1,421 transcripts downregulated by HFD in both age groups, was associated with pathways such as metabolism, cytochrome P450, and steroid hormone biosynthesis, echoing the findings from the corresponding whole-liver cluster (Fig. 2C, F). To better assess the similarity of Clusters 1-3 from whole liver and hepatocytes, we analyzed the overlap of DEGs in the corresponding clusters between whole liver and isolated hepatocytes. While many DEGs are unique to one but not the other, a significant number of genes are shared between whole liver and hepatocytes in all 3 clusters (Fig. 2G-I). Overall, these results show that age modifies the liver’s response to HFD, in many cases amplifying the effects of HFD (Clusters 2 and 3), and these effects are mediated in large part by hepatocytes.

**Figure 2.**
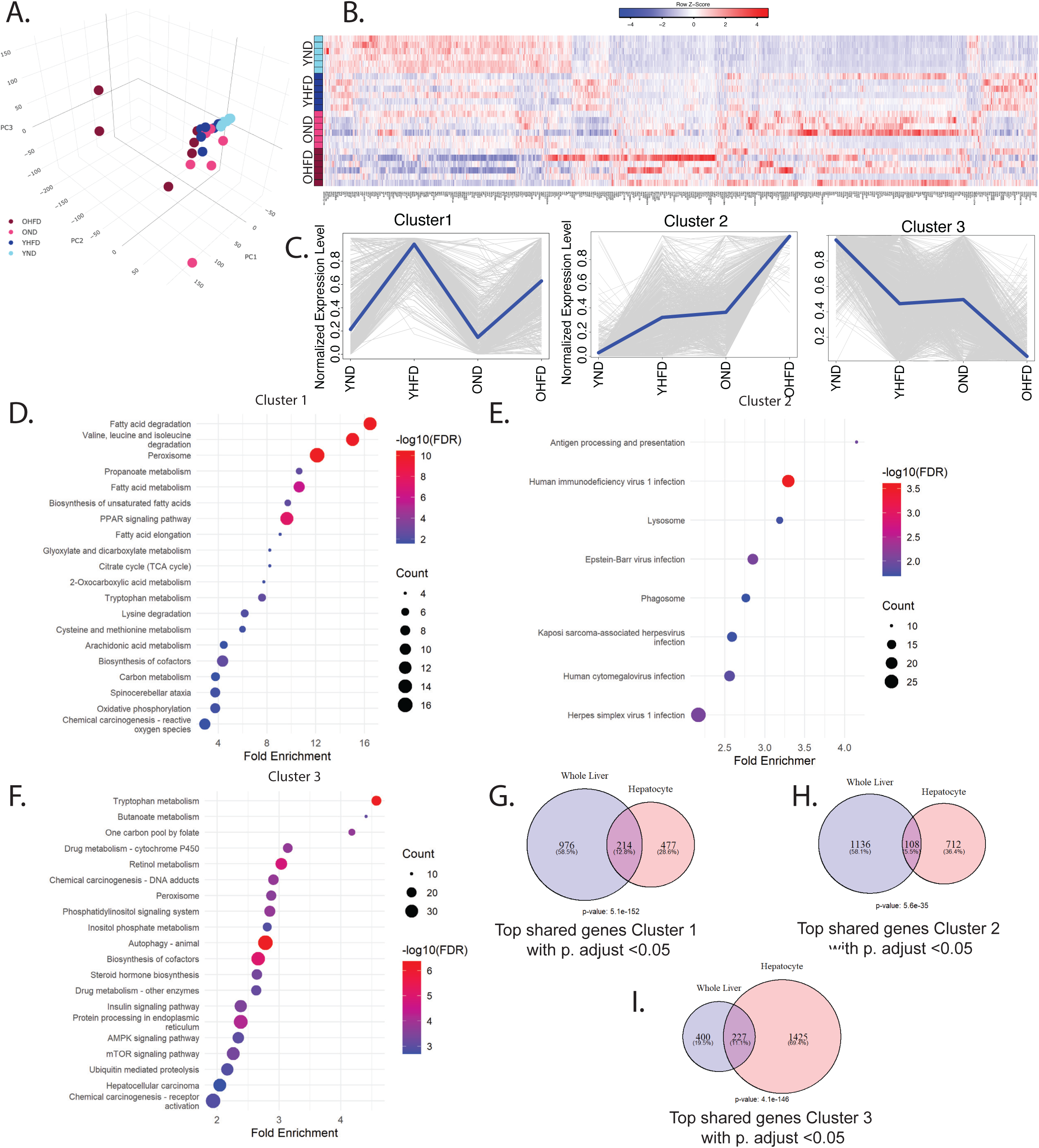
Aged hepatocytes exhibit hyperbolic metabolic dysfunction and heightened proinflammatory signaling in response to HFD. Hepatocytes were isolated from 5-month-old (young) and 22-month-old (aged) mice following 9 weeks of control or high-fat diet (HFD) treatment. (A) Principal component analysis (PCA) of hepatocyte RNA-seq data. (B) Heatmap of all differentially expressed genes (DEGs). (C) Top three gene expression clusters, showing patterns based on age and diet. (D–F) Enriched KEGG pathways corresponding to clusters shown in (C). (G–I) Venn diagrams comparing genes from whole liver and isolated hepatocyte datasets within analogous clusters 1–3, respectively.

Given the interaction between age and HFD, we wondered whether the pro-longevity intervention rapamycin would mitigate the effects of a high-fat diet (HFD) in aged mice. To address this, we administered rapamycin (42 ppm encapsulated in eudragit) or vehicle control to C57BL/6 male mice, beginning at 4 months of age to avoid developmental effects. At 18 months, mice were either maintained on ND or switched to HFD ± rapamycin for 9 weeks, resulting in three cohorts: ND + vehicle, HFD + vehicle, and HFD + rapamycin (Fig. 3A). At 21 months, whole livers were collected for histological analysis, and hepatocytes were isolated for biomarker analysis and RNA sequencing. Western blot analysis of the ratio of phosphorylated S6 to total S6, a marker of mTOR activity, was significantly reduced in hepatocytes of the HFD + Rapa group as compared to the HFD + vehicle cohort indicating on target activity of rapamycin (Fig. 3B-C). PCA of hepatocyte transcriptomes revealed distinct clustering of cohorts, with the greatest number of DEGs observed between HFD + Rapa and HFD + vehicle (Fig. 3D-E). Of note, HFD + Rapa vs ND + Veh showed fewer DEGs than HFD + Veh vs ND + Veh, consistent with an ability of rapamycin to antagonize the effects of HFD (Fig. 1E). Indeed, a plot of all genes differentially expressed between HFD + Veh vs ND + Veh against HFD + Rapa vs HFD + Veh indicated that rapamycin reversed or suppressed most transcriptional changes induced by HFD, with a Pearson correlation of −0.92 between DEGs modulated by HFD and averted by Rapa (Fig. 3F). This is visualized by a heatmap of all genes upregulated in HFD + Veh vs ND + Veh showing a strong suppression by Rapa (Fig. 3G).

**Figure 3.**
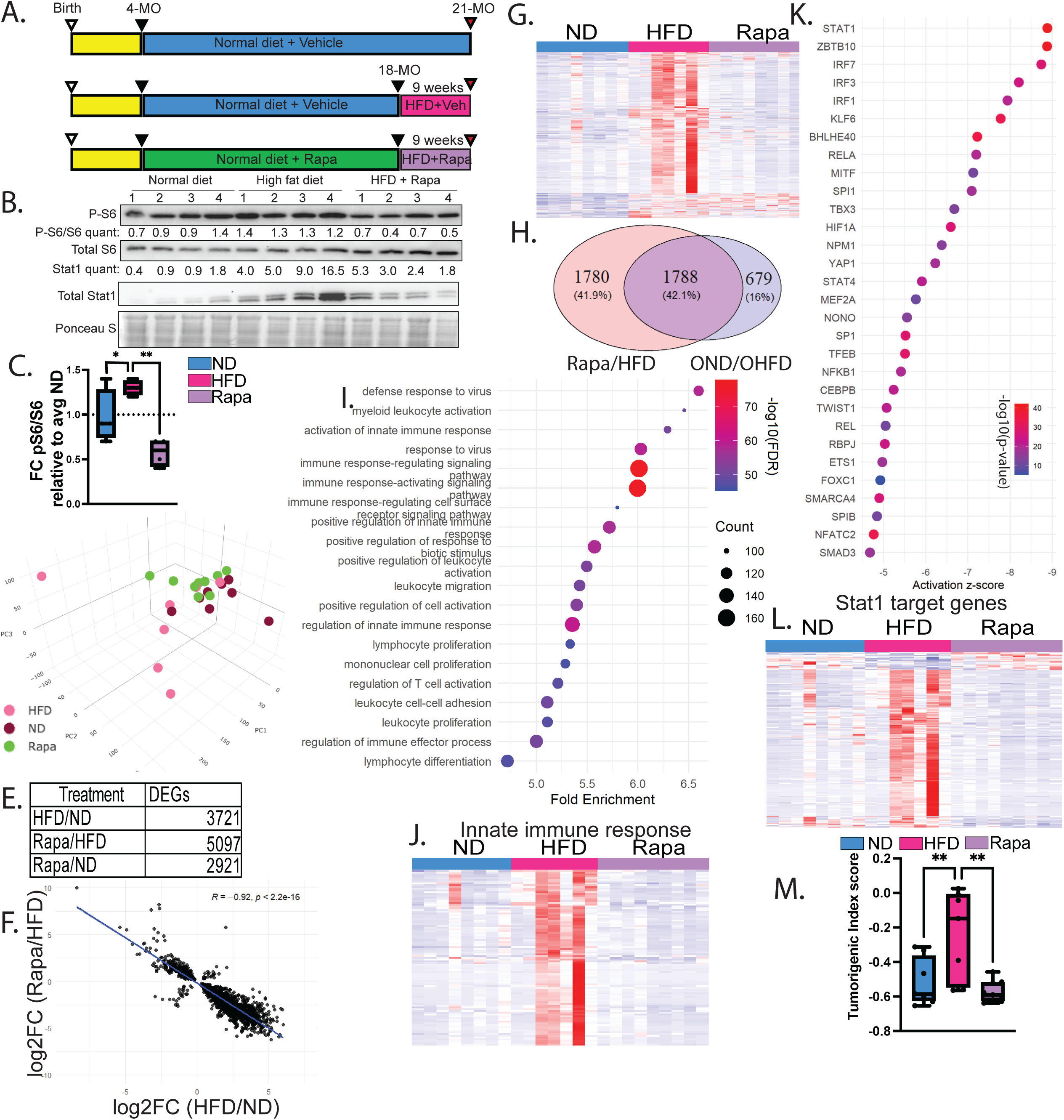
Rapamycin co-treatment mitigates proinflammatory transcriptional hyperactivation in aged mice exposed to HFD. (A) Schematic of treatment protocol: mice received control or rapamycin-containing diet from 4 to 18 months of age, followed by 9 weeks on HFD or continued normal diet. (B) Western blot analysis of phospho-S6 (p-S6), total S6, and Stat1 in liver tissue; Ponceau S used as a loading control. Quantification of p-S6/S6 ratio and Stat1 protein levels shown. (C) Relative fold change of p-S6/S6 compared to control diet. (D) PCA of RNA-seq from isolated hepatocytes. (E) Table summarizing the number of differentially expressed genes (DEGs) across treatment groups. (F) Scatter plot comparing gene expression changes induced by HFD (HFD + Veh vs ND + Veh) with those reversed or suppressed by rapamycin (Rapa + HFD vs HFD + Veh). (G) Heatmap of genes upregulated by HFD + Vehicle treatment. (H) Venn diagram showing overlap between transcripts elevated by HFD versus normal diet and those decreased by rapamycin. (I) KEGG pathway enrichment analysis of overlapping genes. (J) Heatmap of innate immune response gene expression. (K) Top transcription factor pathways predicted by IPA in the overlapping gene set. (L) Heatmap of Stat1 target gene expression. (M) Tumorigenic index score calculated per mouse. Statistical analyses for p-S6/S6 ratios and tumorigenic index scores were performed using one-way ANOVA with post hoc Tukey’s test; *p* < 0.05 (*), *p* < 0.1 (**).

Previous analysis of whole liver and purified hepatocytes identified clusters 1 and 2 as upregulated by HFD (Fig. 2). We asked whether these genes are suppressed by rapamycin. Of 2467 genes upregulated by HFD in hepatocytes of old mice, 72% (1788 genes) were downregulated by rapamycin, a highly significant 6.5-fold enrichment over random (Fig. 3H). These genes were enriched for immune-related processes such as defense response to infection and innate immune activation (Fig. 3I), similar to Cluster 2 induced by HFD (Figs. 1 and 2). A heatmap of the genes associated with regulation of innate immune response shows the effect of rapamycin to assuage this HFD-induced activation of this pathway (Fig. 3J). Ingenuity Pathway Analysis (IPA) highlighted key pro-inflammatory transcriptional regulators, including *Stat1*, *Irf3*, and *Rela*, which were downregulated by rapamycin (Fig. 3K). A heatmap of *Stat1* target genes exemplifies this repressive effect of rapamycin (Fig. 3L). Western blot analysis confirmed elevated *Stat1* protein levels in HFD + vehicle and a marked reduction in rapamycin + HFD (Fig. 3B). To assess the association of these transcriptional changes on risk of cancer, a disease whose incidence is increased with age and metabolic stress, we applied the tumorigenic index (TI) algorithm developed by Wang et al., which calculates the likelihood of hepatocellular carcinoma (HCC) development based on RNA-seq data. The TI, elevated by HFD compared to ND, was completely normalized in the HFD + Rapa cohort (Fig. 3M). Collectively, these results demonstrate that rapamycin suppresses many of the HFD-induced pro-inflammatory and pro-tumorigenic transcriptional changes in livers of aged mice on HFD, indicating a protective effect against inflammation and associated oncogenic stress in the context of aging.

Analysis of whole liver and purified hepatocytes identified a cluster of genes (Figs. 1 and 2, Cluster 3) that was down regulated by HFD in both young and old, but especially in old. Of these 1254 genes down regulated by HFD, 477 were rescued by rapamycin treatment of old mice (p value: 3.49e-88 and Fold Enrichment: 6.6) (Figure 4A). Analysis of the 477 genes downregulated by HFD and upregulated by rapamycin revealed enrichment in metabolic pathways, including cholesterol, icosanoid, sterol, and fatty acid metabolism, as well as protein folding and degradation processes such as ER-associated protein degradation (ERAD) and the ER stress response (Fig. 4B). Pathways conserved from this analysis compared to Cluster 3 in Figure 2 (i.e., downregulated by HFD and upregulated by rapamycin) include ERAD pathway, Olefinic compound metabolic processes, response to ER stress, steroid metabolic process, alcoholic metabolic processes and fatty acid metabolic processes. Heatmaps for fatty acid metabolic processes and response to ER stress demonstrated that these pathways in HFD + rapamycin-treated mice closely resembled those in the normal diet (ND+Veh) group (Fig. 4C-D). Ingenuity Pathway Analysis (IPA) identified key transcriptional regulators of these genes, including *Trim24*, *Sirt1*, and *Hnf4a*, which play important roles in liver metabolic homeostasis and the modulation of inflammation (Fig. 4E)^29–31^. Collectively, these results further show that rapamycin suppresses additional HFD-induced transcriptomic changes in livers of aged mice, indicating a protective effect against metabolic dysfunction in aging.

**Figure 4.**
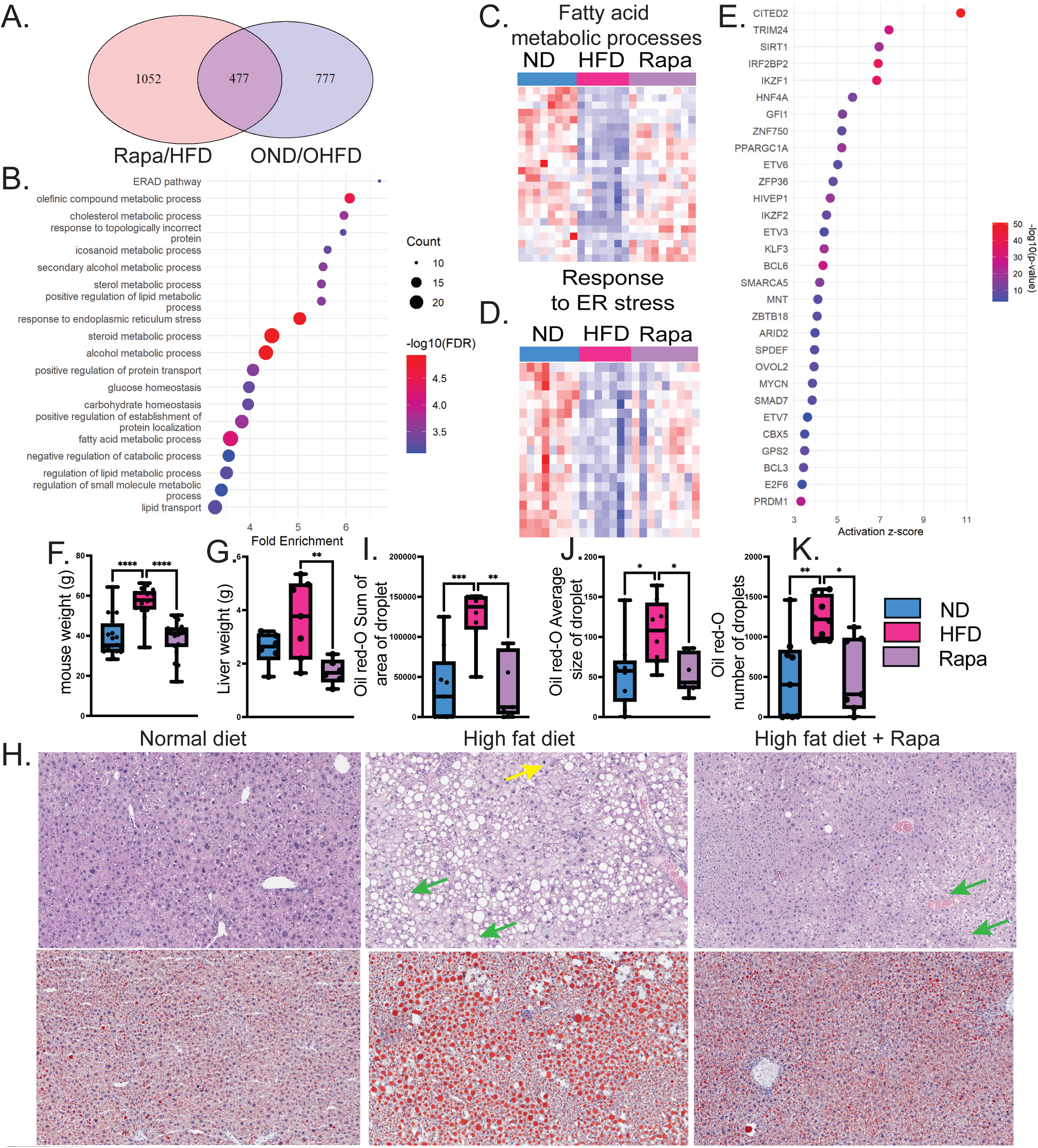
Rapamycin co-treatment restores metabolic pathways prevents weight gain and steatosis in aged mice. (A) Venn diagram of overlapping genes downregulated by HFD vs ND and upregulated by rapamycin; p = 3.49e–88, fold enrichment = 6.615. (B) KEGG enrichment analysis of the 477 overlapping genes. (C) Heatmap of genes involved in fatty acid metabolic processes. (D) Heatmap of genes associated with ER stress response. (E) Ingenuity Pathway Analysis identifying key transcriptional regulators of the 477genes. (F, G) Body weight and liver weight (g) of mice. (H) Representative liver histology: H&E staining (top row) and Oil Red O staining (bottom row). Yellow arrows indicate hepatocyte ballooning; green arrows indicate lipid droplets. (I–K) Quantification of Oil Red O staining: total lipid area (I), average droplet size (J), and droplet number (K). Statistical analyses for weights and Oil Red O quantifications using one-way ANOVA with post hoc Tukey’s test. Significance is denoted as follows: p < 0.05 (*), p < 0.1 (**), p < 0.001 (***), p < 0.0001 (****).

Mice on HFD + vehicle exhibited significant weight gain, with a median final weight of 57.8 g compared to 35.3 g in the ND + vehicle group. In contrast, mice on HFD + rapamycin had a reduced median final weight of 41.3 g (Fig. 4F). Similarly, liver weights were highest in the HFD + vehicle group (3.7 g) compared to ND + vehicle (2.6 g), while HFD + rapamycin showed reduced liver weight to 1.7 g (Fig. 4G). Histological analysis with H&E and Oil Red O staining showed extensive lipid accumulation (Fig. 4H, green arrows) and hepatocyte ballooning (Fig. 4H, yellow arrows) in the livers of HFD + vehicle mice, whereas the addition of rapamycin markedly reduced lipid deposition, restoring liver morphology closer to that of the ND + Veh group (Fig. 4H-K). These findings demonstrate that rapamycin is able to assuage the response to metabolic stress in old mice, as reflected in normalization of metabolic transcriptional profiles and prevention of inflammation-associated signaling, a reduction of total body weight and suppression of features of steatotic liver disease and NASH.

## Discussion

This study demonstrates that aging amplifies components of the hepatic response to HFD-induced metabolic stress and that rapamycin treatment effectively mitigates these HFD-induced transcriptional and physiological changes in aged liver. Whole liver and hepatocyte-specific RNA-sequencing revealed that aging exacerbates the upregulation of pro-inflammatory signaling pathways by HFD, while promoting the downregulation of genes involved in metabolism, cytochrome p450 and steroid hormone biosynthesis. Rapamycin treatment of aged HFD-fed mice reversed many of these changes, including inflammatory transcriptional programs (e.g., *Stat1*, *Rela*) and suppression of metabolic expression. Together, these findings position aging as a key modifier of liver susceptibility to dietary metabolic stress and highlight rapamycin as a promising intervention to restore metabolic homeostasis and prevent diet-associated disease progression in the setting of old age.

Aging is associated with increased vulnerability to metabolic stress, which contributes to disproportionate risk for metabolic-associated fatty liver disease (MAFLD) and related health disparities in older populations. Older adults experience physiologic changes that impair metabolic flexibility, including reduced mitochondrial efficiency, impaired insulin signaling, and chronic low-grade inflammation^28,32,33^. These age-related alterations heighten susceptibility to lipid accumulation, hepatic steatosis, and systemic metabolic dysfunction when exposed to a sedentary lifestyle, high-fat diets or other metabolic stressors^25,34–38^. Moreover, aging is accompanied by changes in liver cell composition and function, including hepatocyte senescence and increased immune cell infiltration, which may accelerate MAFLD progression^4,18,39–41^. Clinical evidence indicates that older individuals are more likely to experience severe steatosis, fibrosis, and adverse outcomes associated with MAFLD, yet are often underrepresented in therapeutic trials^24,42,43^. Understanding how age alters the liver’s response to metabolic stress is critical for developing interventions that address the specific vulnerabilities of older individuals.

With age and metabolic stress, the cellular composition of the liver shifts, immune infiltration increases, hepatocyte number and function decline, and fibrosis may alter the representation of different cell types, confounding interpretation of whole-tissue transcriptomes^4,24–28^. Isolating hepatocytes enables dissection of their autonomous transcriptional responses, distinguishing direct effects of aging and HFD on hepatocytes from effects secondary to changes in tissue composition. Hepatocytes play a central physiological role in the progression of metabolic dysfunction, serving as the primary site of lipid accumulation, insulin responsiveness, ER stress, and early inflammation in MAFLD. Within this study, hepatocyte-specific transcriptional programs regulating metabolism and immune signaling were more clearly resolved when analyzed independently from the complex whole-liver transcriptome.

Rapamycin has emerged as a promising intervention capable of simultaneously targeting age-related decline and maladaptive responses to metabolic stress. Through inhibition of mTORC1, rapamycin suppresses anabolic pathways and promotes a shift toward oxidative metabolism and mitochondrial efficiency^44–46^. It enhances fatty acid oxidation and activates transcriptional regulators like *Ppar*α and *Sirt1*, supporting improved lipid metabolism and stress resilience. Rapamycin has been shown to restore metabolic gene expression, reduce adiposity, and alleviate hepatic steatosis in aging and metabolic disease models^21,47,48^. While the benefits of rapamycin have been characterized in either metabolic disease or aging models individually, they have not been thoroughly explored in the context of their intersection, where aging amplifies vulnerability to metabolic stress. Our findings position rapamycin as a therapeutic candidate capable of restoring metabolic resilience in the aged liver, offering a strategy to counteract the compounding effects of age and dietary stress on metabolic health.

These results suggest that rapamycin can have dual benefit through its ability to promote healthy tissue function and reduce cancer risk. Steatosis is a risk factor for MAFLD progression and liver cancer development. Histological analysis confirmed that rapamycin reduced liver steatosis and inflammation associated with HFD feeding in aged animals. Interestingly, rapamycin treatment also resulted in a marked reduction in liver tumorigenic index scores in HFD mice (Fig.3M). Our findings align with previous work on rapamycin increasing lifespan and delaying spontaneous tumor incidence in mice^19,49,50^. Thus, rapamycin may serve as a gerotherapeutic that simultaneously promotes metabolic health and cancer resistance in aging.

This study identifies aging as a key enhancer of liver vulnerability to dietary stress and highlights rapamycin as a potent modulator of this response. Aging exacerbates liver transcriptional responses toward metabolic dysfunction, inflammation and tumorigenesis when exposed to HFD. Rapamycin restores metabolic gene expression and suppresses inflammatory programs in aged hepatocytes of HFD fed mice. This intervention reduces liver steatosis, body weight, and tumorigenic risk scores. Findings support broader application of gerotherapeutics in suppressing combined age- and diet-related metabolic dysfunction and cancer risk. These results provide rationale for use of gerotherapeutics as a disease-modifying strategy in diet-induced metabolic disease in older people.

### Experimental Procedures

#### Animal usage

All animal procedures were approved by the Institutional Animal Care and Use Committee (IACUC) of Sanford Burnham Prebys Medical Discovery Institute. Animal experiments were performed at Sanford Burnham Prebys Medical Discovery Institute Animal Facility in compliance with the IACUC guidelines. The studies performed within this manuscript were performed with all relevant ethical regulations regarding animal research. Young and old C57BL/6 animals were obtained from the NIA aging colony housed at Charles Rivers or breed within the institute or were purchased from Charles Rivers. Animals were housed 5 mice per cage and maintained under controlled temperature (22.5 °C) and illumination (12 h dark/light cycle) conditions. High fat diet utilized within this study were obtained from Test Diet 58Y1 consisting of 60% calories from fat or “normal diet” 58Y2, 10% calories from fat. Microencapsulated rapamycin (42ppm) and eudragit control were purchased from Emtora (formerly Rapamycin Holdings) and compounded into appropriate diet by NewCO Distributors. Treatment is similar as described previously^51,52^. Mice were maintained on diet ad libitum and diet was replaced weekly.

#### Hepatocyte isolation

following a protocol adapted from Charni-Natan et al., 2020. Mice were euthanized via CO_₂_ asphyxiation, and once immobilized, the abdominal cavity was opened to expose the inferior vena cava. A 24G × ¾” catheter (Surflo, SR-0X2419CA) was inserted into the vein, and the liver was perfused with 50□mL of pre-warmed (42°C) HBSS (Gibco, Ref #14175-095) at a flow rate of 6□mL/min, followed by 40□mL of DMEM (Gibco, Ref #10313-021) containing Collagenase Type IV (Gibco, Ref #17104-019), also pre-warmed to 42°C. After digestion, livers were excised and gently dissociated in ice-cold DMEM supplemented with 2% Bovine Serum Albumin (BioWorld, CAS: 9048-46-8) and kept on ice. Cells were washed by centrifugation at 50 × g for 2 minutes at 4°C, with media changes repeated five times. The final cell suspension was resuspended in a gradient solution consisting of 40% Percoll (Cytiva, Ref #17089102), 10% 10× HBSS (Gibco, Ref #14065-056), and 60% DMEM, and centrifuged at 100 × g for 7 minutes at 4°C. Viable hepatocytes from the pellet were collected, washed once in DMEM at 50 × g, and counted using a hemocytometer with trypan blue exclusion for viability assessment.

#### Histology and Staining

Picro-Sirius Red staining was performed on 5□µm formalin-fixed paraffin-embedded (FFPE) liver sections following the protocol detailed by Emory University Microscopy in Medicine (see: <u>link</u>). For proliferation analysis, EdU incorporation was carried out using the Click-iT™ EdU Imaging Kit according to the method described by Salic et al. 2008^53^. EdU was administered intraperitoneally at a dose of 140□µL per mouse in PBS, 36 hours prior to euthanasia.

#### Western Blotting

Homogenized tissue or cell suspensions were quantified using the Bradford assay (Pierce, Ref #1863028) on a SpectraMax 190 plate reader. SDS-PAGE gels were freshly cast and run following the protocol described by Lynch et al. (PMID: 27791595) using the Bio-Rad Mini-PROTEAN Tetra System. Proteins were transferred to PVDF membranes (Immobilon-PSQ, Millipore ISEQ00010) at 100 V for 70 minutes. Membranes were stained with Ponceau S (Sigma-Aldrich, P7170-1L) prior to blocking with 5% skim milk (BD Difco, Ref #232100) in TBST for 1 hour at room temperature. Primary antibodies (see Supplementary Table 6) were incubated in 5% BSA (BioWorld, Ref #9048-46-8) overnight at 4°C with rocking. Secondary antibodies used were Goat anti-Mouse IgG-HRP (Thermo Fisher Scientific, Cat# 31446, RRID:AB_228318) and Goat anti-Rabbit IgG-HRP (Millipore, Cat# AP307P, RRID:AB_92641). Membranes were imaged for Ponceau S and HRP signal using a Bio-Rad ChemiDoc Touch system, and images were processed and analyzed with Image Lab software. Antibodies used: total Stat1 (Cell Signaling Technologies 9172S), S6 Ribosomal protein (Cell Signaling Technologies 2317), phospho-S6 Ribosomal protein (Cell Signaling Technologies 5364).

#### RNAseq Analysis

Raw fastq files were aligned to mm10(Gencode vM23), using STAR^54^ 2-pass pipeline. Aligned reads were filtered, sorted, and indexed by SAMtools V1.1.0^55^. Genome tracks (bigWig files) were obtained by Deeptools V3.3.2^56^. Raw read counts were obtained by HTSeq V0.11.2 for differential analysis. Differentially expressed genes were obtained by DESeq2 V1.26.0^57^ CuffLinks V2.2.1Trapnell^58^ was used to compute FPKM values.

#### Clustering and Visualization of Differentially Expressed Genes

Differentially expressed genes (DEGs) were clustered based on their normalized FPKM expression values using hierarchical clustering with Pearson correlation distance and average linkage. Genes with missing or zero expression were removed, and expression values were scaled by row. Clusters were defined by cutting the dendrogram into groups **f**or each gene cluster, normalized expression profiles were visualized by min-max scaling each gene’s expression across samples and plotting both individual gene trajectories and the average expression profile. These plots were used to assess shared expression dynamics across experimental groups.

#### Principal Component Analysis

(PCA) was performed on log_₂_-transformed FPKM values (with a pseudocount of 1) after transposing the matrix to set samples as rows. Genes with zero variance were excluded. PCA was conducted using the prcomp function in R, and the first three principal components were visualized using an interactive 3D scatter plot generated with the plotly package. Samples were colored by group to assess separation based on expression profiles.

#### Gene Set Overlap and Functional Enrichment Analysis

Overlapping up- and downregulated genes from two differential expression comparisons were identified and visualized using Venn diagrams (VennDiagram), with overlap significance calculated via hypergeometric testing and high-precision p-values (Rmpfr). Fold enrichment was computed using a mouse genome background of 32,179 genes.

#### GO and KEGG enrichment analyses

were conducted with clusterProfiler (q < 0.05, BH correction), and top terms were visualized using dot plots showing fold enrichment and –log_₁₀_ adjusted p-values. For selected GO terms, Z-score–normalized expression values were used to generate heatmaps of associated genes across experimental groups.

#### Ingenuity Pathway Analysis (IPA)

Upstream regulator analysis was performed using Ingenuity Pathway Analysis (IPA) and filtered to include only transcription factors. Transcription factors with significant overlap (*p* < 0.05) and non-zero activation z-scores were retained. The top 30 predicted activators and inhibitors were ranked by activation z-score and visualized as dot plots, with color indicating –log_₁₀_ *p*-value. Positive z-scores reflect predicted activation, while negative z-scores indicate predicted inhibition.

#### Correlation Analysis of Gene Expression Changes

To assess how Rapamycin modulates gene expression changes induced by a high-fat diet (HFD), log_₂_ fold changes (log_₂_FC) were calculated for ND vs HFD and HFD vs Rapamycin using normalized FPKM values. The analysis included genes that were differentially expressed in the HFD condition relative to ND. Log_₂_FC values were computed using group means and a pseudocount (ε = 1e–3) to avoid division by zero. A scatter plot was generated to compare the two contrasts, and Pearson correlation was used to quantify their relationship.

#### Use of artificial intelligence technology

ChatGPT was used for grammatical and formatting purposes with human oversight. The ideas and development of the manuscript along with results and conclusions were generated and reviewed by authors on the manuscript prior and post writing.

